# Flexible color segmentation of biological images with the R package recolorize

**DOI:** 10.1101/2022.04.03.486906

**Authors:** Hannah I. Weller, Steven M. Van Belleghem, Anna E. Hiller, Nathan P. Lord

## Abstract

Color is an important source of biological information in fields ranging from disease ecology to sexual selection. Despite its importance, most metrics for color are restricted to point measurements. Methods for moving beyond point measurements rely on color maps, where every pixel in an image is assigned to one of a set of discrete color classes (color segmentation). Manual methods for color segmentation are slow and subjective, while existing automated methods often fail due to biological variation in pattern, technical variation in images, and poor scalability for batch clustering. As a result, color segmentation is the common bottleneck step for a majority of existing downstream analyses. Here we present `recolorize`, an R package for color segmentation that succeeds in many cases where existing methods fail. `Recolorize` has three major components: (1) an effective two-part clustering algorithm where color distributions are binned and combined according to perceived similarity in a frequency-independent manner; (2) a toolkit for minor manual adjustments to automatic output where needed; and (3) flexible export options. This paper illustrates how to use `recolorize` and compares it to existing methods, including examples where we segment formerly intractable images, and demonstrates the downstream use of methods that rely on color maps.

## Introduction

Color is an important source of biological variation, and is an important trait in a wide range of biological questions, ranging from sexual selection, camouflage, and animal communication, to thermal physiology, disease ecology, development, and genetics (Bekker et al., 1837; Bates, 1863; Poulton, 1890; Orteu and Jiggins, 2020; Hooper et al., 2020; Van Belleghem et al., 2020). Despite their obvious importance in biology (e.g., signal function, taxon identification, color production and variation), there is little consensus about how to quantify and compare color patterns, even though this is a necessary first step in testing most questions about them. Contrast this with, for example, geometric morphometrics, a family of methods for quantifying variation in biological shape (Bookstein, 1996). Researchers have reached a general consensus about how to quantify and compare morphology (Klingenberg, 2011; Adams and Otárola-Castillo, 2013; Olsen and Westneat, 2015). This consensus, coupled with the availability of a handful of software tools for easily digitizing specimens, has led to a marked increase in both the number of studies that use geometric morphometrics and the insights resulting from this work (Polly et al., 2013; Adams et al., 2004; Lawing and Polly, 2010). There is no such set of consensus methods for measuring variation in biological color.

Partly, lack of consensus is inevitable—color patterns are multidimensional and receiver-dependent, so variation can come from a wide range of biological and technical sources (this true to some extent for morphology, but at least this variation all exists in the same three dimensions). Individual organisms might vary in the location, arrangement, and intensity of their color pattern elements; the optical physical properties of those patterns (pigmentary or structural); and the surface topology and 3D morphology of the organism itself. Even if these sources of variation are well-defined, the ways in which we detect and record this information can introduce noise and error.

Color pattern data is usually measured from images captured through digital sensors constructed from human-centric visual system sensitivities, which can (and do) vary from camera to camera. The colors reflected by a surface also depend on the available light, whether natural or artificial, which varies in emission output, intensity, and direction, all of which will affect how light reflects off of a given surface (with its own material properties) (Johnsen, 2012). Given these constraints, it is not surprising that universal methods for measuring color patterns in a consistent and repeatable way have remained elusive. How do we meaningfully measure the difference between organisms that may not have homologous color pattern elements? What if the images were taken with different lighting conditions and cameras? What do we compare? How much can we reduce this dimensionality and still retain the relevant variation—assuming we even know what constitutes relevant variation?

Plenty of available methods exist for measuring particular aspects of color patterns, but most of these are specific to a system or question. Researchers often have no choice but to build a new method specific to their problem (Caves and Johnsen, 2018; Valcu and Dale, 2014; Gawryszewski, 2018; Chan et al., 2019; van den Berg et al., 2020; Troscianko et al., 2017; Yang et al., 2016; Van Belleghem et al., 2018; Maia et al., 2019; Weller and Westneat, 2019; Hooper et al., 2020; Valvo et al., 2021; Endler et al., 2018; Endler, 2012; Ezray et al., 2019). This sets an unreasonably high barrier to entry for measuring color patterns: developing methods is time-consuming and measurements are difficult to compare.

Some of the most frequently used and promising methods for quantifying color pattern variation are also the most flexible, in that they can be applied to a wide variety of organisms and pattern types. Well-known examples include Endler’s adjacency and boundary strength metrics (Endler, 2012; Endler et al., 2018), which emphasize contrast between adjacent color patches; the `patternize` package (Van Belleghem et al., 2018), which quantifies color pattern variation essentially using a sampling grid); the micaToolbox suite of tools for multispectral image analysis (van den Berg et al., 2020); and the matching of spatial image data with spectral reflectance data in the `pavo` package (Maia et al., 2019). Most of these metrics require first clustering the image into discrete color patches (making “color maps”) and then measuring how various aspects of these patches vary across images (Fig. 1). Essentially, they start from the simplifying assumption that a color pattern can be discretized into regions of uniform color, and then compare the shape and color of those regions in some biologically relevant way.

**Figure 1:**
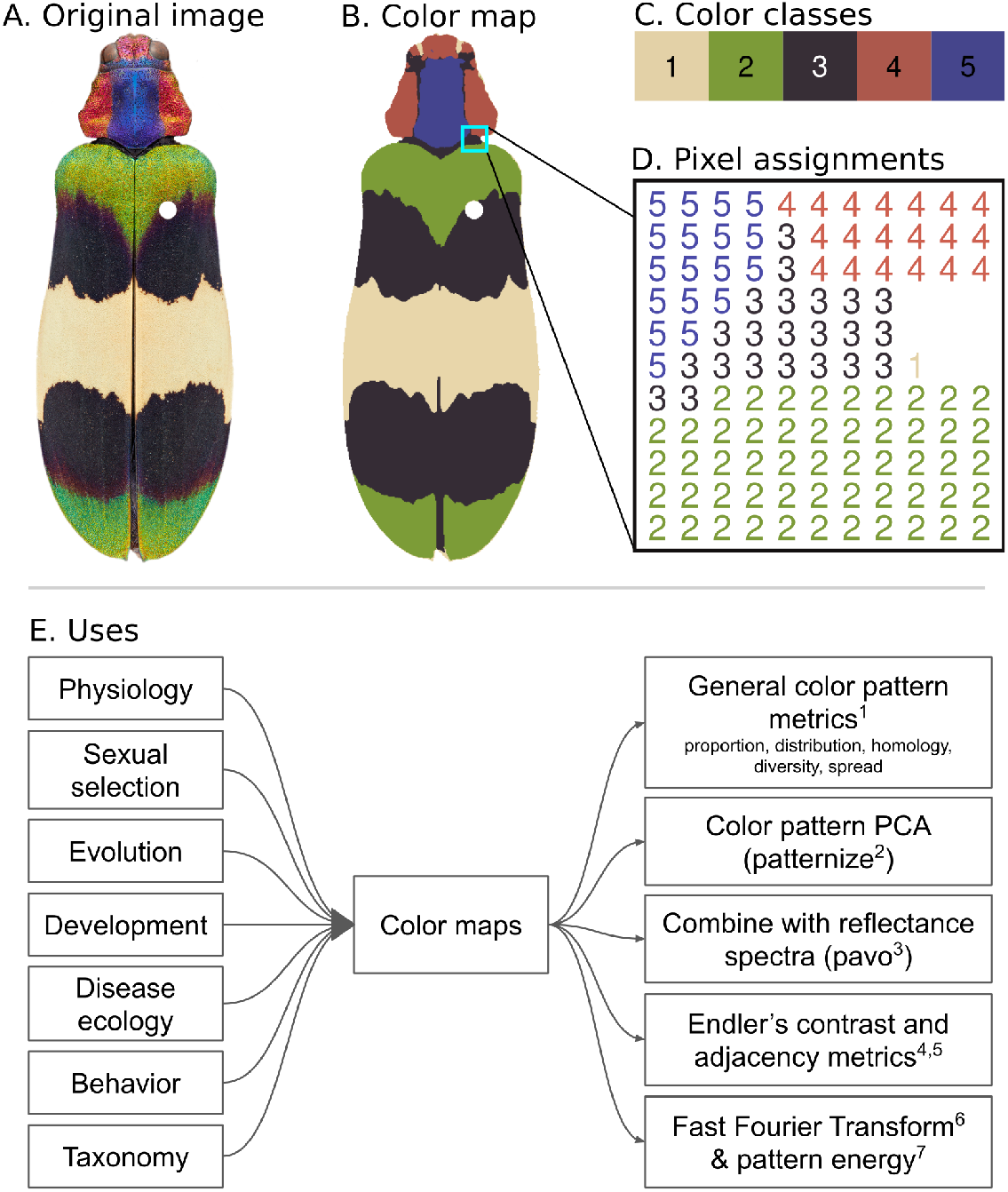
Color map example made with `recolorize`. A: Original image of a beetle, *Chrysochroa corbetti*. Limbs, antennae, and insect pin have all been masked. B: Color map, where each pixel has been assigned to one of five color classes. C: The five color classes displayed as a color palette; in this case the displayed colors of the palette (the color centers) represent the average red-green-blue (RGB) color of all pixels assigned to that class. D: A representative section of the color map as a numeric matrix: each pixel is assigned to a color class, so that we can refer to the entirety of color patch 2 by indexing all values of the color map equal to 2, which is associated with a particular color (in this case, green). E: Color maps are a major bottleneck step of color pattern quantification. The boxes on the left list some fields for which color pattern is an important trait; on the right are commonly used metrics or methods for quantifying color pattern, all of which require color maps as a starting point. Original image: Nathan P. Lord. Citations: 1. Chan et al. (2019); 2. Van Belleghem et al. (2018); 3. Maia et al. (2019); 4. Endler (2012); 5. Endler et al. (2018); 6. Mason et al. (2021); 7. van den Berg et al. (2020).

Generating color maps from digital images–in a reproducible and efficient way–has proven to be the bottleneck step for these methods, especially across sets of images (Fig. 1E). Image segmentation in general is a notoriously complex problem, compounded here by the fact that the number and boundaries of color patches are often ambiguous; there isn’t necessarily a ‘correct’ answer, just one appropriate to the biological question. An image without some kind of segmentation, however, is little more than a pile of pixels; segmentation provides labels to groups of pixels so that we can refer to (and measure) particular regions. Researchers typically choose between automated methods, which require little or no user input but are difficult to modify when they do not work well, and manual segmentation, which is typically slow and subjective (Hooper et al., 2020).

The most widely used automated method is k-means clustering (Hartigan and Wong, 1979), which has been implemented in several R packages for color pattern analysis (Weller and Westneat, 2019; Van Belleghem et al., 2018; Maia et al., 2019), making it more accessible than other approaches. Users specify only the expected number of color classes, and the k-means algorithm attempts to find the set of color classes that minimize within-cluster variances. While this is a relatively intuitive and flexible approach in theory, in practice k-means clustering suffers from a number of issues:

1. In trying to minimize variances, k-means tends to over-cluster large color patches and fails to differentiate small, colorful patches, meaning it often misses details and is highly sensitive to shadows, 3D contours, texture, and specular reflections (all frequent features of organism photographs) (Fig. 2A-B).
2. Users can’t compare images that were taken from different sources or under different lighting conditions, because they can produce very different color distributions (Fig. 2C) that k-means clustering can’t correct for.
3. Users have to specify the number of color classes, which is subjective, and for comparative datasets may not be the same for all species (Fig. 2D).
4. The implementation of the algorithm is usually heuristic, not deterministic. The same image will result in different color clusters depending on the run, and the color classes themselves are returned in an arbitrary order. This makes it difficult to perform batch processing when we want to map a set of images to the same set of colors, because yellow might be color 1 in our first image and color 2 in our second image (Fig. 2E), and the colors will be in a new random order every time we rerun the analysis unless users set a particular seed at the start of a clustering session.

**Figure 2:**
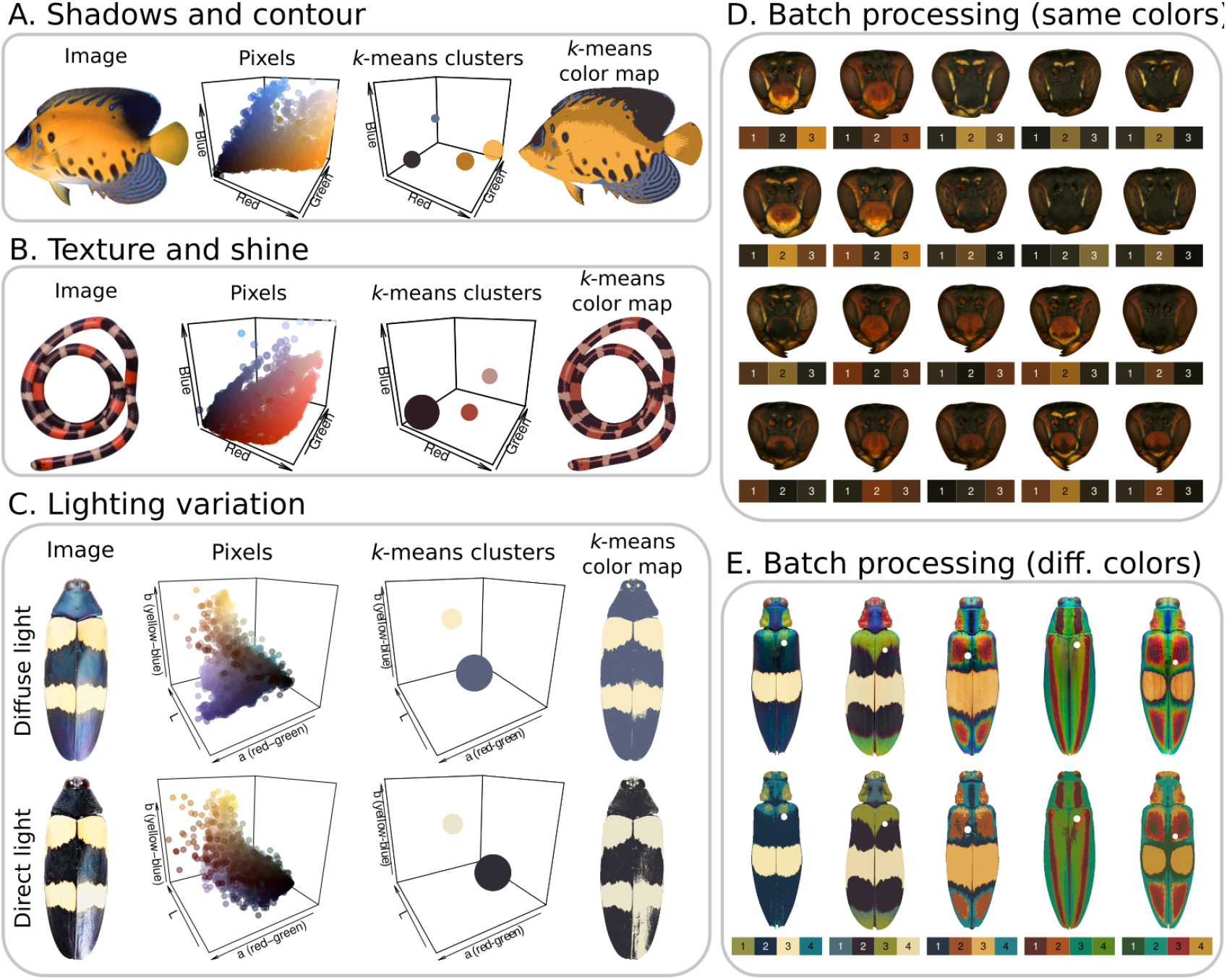
Examples of images and image sets for which k-means clustering is inadequate for color segmentation. A: This angelfish (*Pygoplites diacanthus*) has four color classes: yellow, black, blue, and white. K-means achieves lower within-cluster variances by assigning a light and dark yellow color center when we fit n = 4 centers. B: A coral snake (*Micrurus* sp.) with three color classes: red, white, and black. Specular highlights (shine) and scale texture result in shiny areas of the snake being assigned to the white color center. C: The same beetle specimen (*Chrysochroa mniszechii*) photographed using diffuse (upper row) and direct (lower row) light. Fitting n = 2 color centers using k-means clustering produces different color centers and different color patch geometries. D: Batch processing with the same set of color classes. All of these *Polistes fuscatus* wasp faces share a palette of three colors (yellow, reddish brown, and dark brown), but fitting n=3 color centers for each image using k-means clustering results in subtly different colors for each image, and they are returned in an arbitrary order. E: Batch processing for image sets with different types and numbers of colors. We chose to fit n = 4 colors for each beetle (*Chrysochroa* spp.) in this 5-species dataset, resulting in some images being over-clustered and some being under-clustered. Image sources: Jack Randall (A), Alison Davis-Rabosky (B), Nathan P. Lord (C & E), James Tumulty (D).

There are other methods for performing color segmentation, but they share many of the same weaknesses as k-means or are otherwise too limited in scope to be as easily generalized to a range of problems. Edge detection methods (e.g. watershedding, Otsu algorithm, Canny operator) use spatial information, which can resolve glare and noise to some extent, but only work well for segmenting color patches with sharp boundaries and often treat textures like scales as edges. The receptor-noise limited (RNL) clustering implemented in the Quantitative Color Pattern Analysis toolbox for ImageJ (van den Berg et al., 2020), which uses properties of the visual system to segment colors based on perceived color differences, works well under specific assumptions. However, it addresses only a subset of biological questions about color pattern variation, and as a result has more stringent requirements for data and equipment (and thus a higher barrier to entry). We could also find no dedicated tools specifically for making color maps: most methods provide a function or functions for performing color segmentation before running the rest of the analysis, so users who need color maps for any other purpose not yet implemented in these tools need sufficient coding expertise to extract them.

The color segmentation problem has proven to be so intractable that quantitative color pattern analysis has mostly been limited to organisms with brightly colored, highly contrasting color patterns on relatively flat and untextured surfaces. This largely limits us to some species of butterflies and poison frogs. As a result, our ability to generate questions and hypotheses about color pattern evolution have far outpaced our ability to quantitatively analyze them.

The `recolorize` package is designed to address this problem so that downstream color pattern analysis tools can be used for a wider variety of contexts. We aim to make the package easy to use, easy to modify, and easy to export to other packages and pipelines. The color segmentation options are fast and deterministic, have been tested across a wide variety of images, organisms, and use cases, and are reasonably straightforward in their structure. The general process is: 1. initial clustering step; 2. refinement step; 3. optional semi-manual edits; 4. export to desired format. The resulting toolbox is capable of handling a wide range of images, and has been used successfully in several contexts where k-means clustering has not worked. In order to illustrate that functionality, this paper will go over how to use `recolorize` to generate usable color maps for all of the examples which fail with k-means clustering in Fig. 2.

## Glossary

Terms used for each of the components of this process vary somewhat in the literature; here we define what we mean in this paper (and in the package) for reference.

1. **Color pattern**: Static visual appearance of the sender, e.g. how colors are arranged on the organism, as captured by a single image (or 3D surface).
2. **Color class**: A specific ID (usually numeric) to which portions of the color pattern are assigned.
3. **Color center**: The computer-readable color of the color class, typically expressed as an RGB triplet.
4. **Color patch**: All the portions of a color pattern which are assigned to the same color class.
5. **Color space**: The coordinate system (typically three-dimensional) used to represent the color of each pixel, and within which color distances are calculated.
6. **Color segmentation**: The process of segmenting an image into discrete color patches.

## Results

Code and images for running each of these examples is provided at https://github.com/hiweller/recolorize_examples. All examples are deterministic, meaning that the code as written will produce the same results every time it is run, regardless of the R seed or computer.

### Preparing images for recolorize

Many users of `recolorize` will already have images, and the package is fairly forgiving of variation due to lighting, texture, or arrangement (e.g., of feathers on a bird) since we include tools for making post-hoc adjustments. In brief, users should try to control for as many sources of variation as possible: use the same camera, lighting, background, resolution, positioning, and color standard for the entire dataset.

The most important pre-processing step for using `recolorize` is background masking, for which there are many existing tools available. The package does not perform background masking or image segmentation: instead, users should mask out the background of the image using transparencies (e.g. using GIMP or Photoshop). `Recolorize` can also ignore a background of uniform color by specifying a range of RGB colors. There are several tools available for automatic background masking, such as Sashimi (Schwartz and Alfaro, 2021) or Batch-Mask (Curlis et al., 2021), which could be used for large image sets. The exception is for the `patternize` workflows we show in examples E and F below, where the landmark alignment performs automatic background masking on unmasked images.

### Examples

#### Example A: Main recolorize workflow

The major problem in Fig. 2A is that by fitting *n* = 4 color centers for the angelfish image, k-means clustering achieves lower within-cluster variances by assigning separate light and dark yellow color centers (since these take up a greater proportion of the image) and grouping the white pixels into the light yellow color class, despite the fact that these colors are less similar than the dark and light yellow (as indicated by their proximity in 3D color space). This is a common issue with k-means clustering that is easily resolved by the basic `recolorize` workflow, so we will use this image to illustrate these steps in detail.

The core of the `recolorize` package is a two-step process for color segmentation: first, each pixel in the image is assigned to a region based on ranges defined for each color channel, resulting in what is essentially a 3D histogram of color distribution for the image (Fig. 3A). This is accomplished with the `recolorize` function, using a user-specified number of bins per channel, where the total number of resulting color centers will be n3 for n bins per channel (here, we used 3 bins per channel resulting 3^3^ = 27 possible color regions, only 9 of which contained pixels from the fish image). The color center of each region is then calculated as the average value of all of the pixels assigned to that region (or the geometric center if no pixels were assigned to it). This step requires no distance calculations or color center estimations because pixels are binned into pre-existing regions. As a result, this step is relatively fast and deterministic, and serves to reduce potentially millions of colors in the original image to typically only a few dozen, depending on the user’s specification of how finely to bin them.

**Figure 3:**
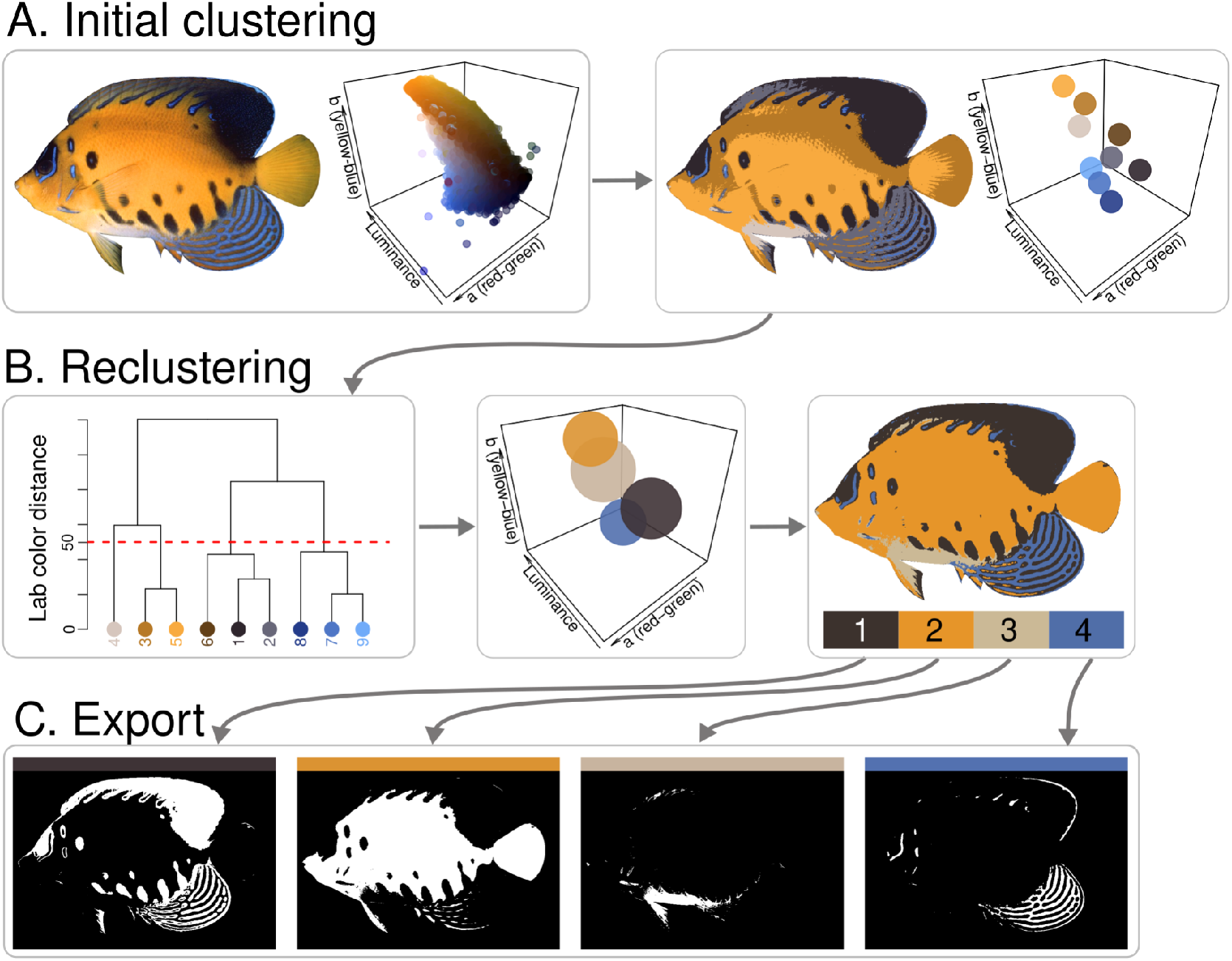
Successful segmentation of the angelfish (*Pygoplites diacanthus*) image from Fig. 2A, illustrating the core steps of the package. A: First, the pixels of the original image are binned by their coordinates in each color channel using a user-selected number of bins per channel using the `recolorize` function. In this case, only 9 of 27 bins had pixels assigned to them. B: These initial bins are combined based on perceived similarity using `recluster` by combining either bins that have a Euclidean distance less than the user-selected cutoff (here, cutoff = 50), or by specifying a final number of colors. The original image is then re-fit using the resulting set of color centers. C: The resulting color map is exported to any of a number of formats. Here, individual color patches are exported as binary masks using the `splitByColor` function.

Second, these initial color centers are reduced according to some rule, such as combining similar color centers or dropping the smallest color patches. The most generally effective function for this step is the `recluster` function, which calculates the Euclidean distance between all pairs of color centers to which any pixels were assigned. Color centers are then clustered by similarity using hierarchical clustering (R Core Team, 2022), and users provide either a similarity cutoff which determines which colors to combine or a final number of expected colors. In Fig. 3B, we used a cutoff of 50 (Euclidean distance in CIE Lab color space) to combine the 9 initial color centers into 4 consensus color centers. New color centers are calculated as the weighted average of the original color centers being combined, with color patch size as the weights. The `recluster` function then refits the original image with these new color centers.

In short, the first step reduces the image to a manageable number of colors, after which more computationally intensive steps (such as calculating a pairwise distance matrix) can be performed deterministically. Because they are also not density dependent, this two-step method also better preserves small but distinct color patches, such as the white patch on the ventrum of the fish image. Finally, the color map can be exported to a variety of formats and packages. In Fig. 3C, we export the color map as a stack of binary masks (0 = color absence, 1 = color presence), but the later examples will illustrate functions for exporting to specific packages.

#### Example B: More complex segmentation

The coral snake from Fig. 2B represents a more complex example, because the specular highlights created by the shiny scales of the snake appear white regardless of their location on the snake (so they cannot be combined with appropriate color centers based on similarity). Instead, we first blur the image to reduce minor color variation due to the scale texture using blurImage, then produce an initial color map using `recolorize` and recluster as in the prior example, resulting in four color centers (Fig. 4). Note that color center 1 (a medium gray) contains most of the specular highlights; remaining portions of the highlight have been assigned to color center 4. This is a fairly common problem in color segmentation output: the method mostly works, but has small problems that render the color map ineffective, with no easy way for users to modify it. For this reason, the `recolorize` package also includes tools for modifying color maps in specific, reproducible ways. First, we can combine color centers 1 and 2 using the `mergeLayers` function. Second, to clean up the small areas of highlight which were assigned to color center 4, we use the `absorbLayer` function: this targets areas of a color patch within a user-specified area and/or location boundary, and changes the color of each separate area to that of the color patch with which it shares the longest border, effectively “absorbing” stray speckles.

**Figure 4:**
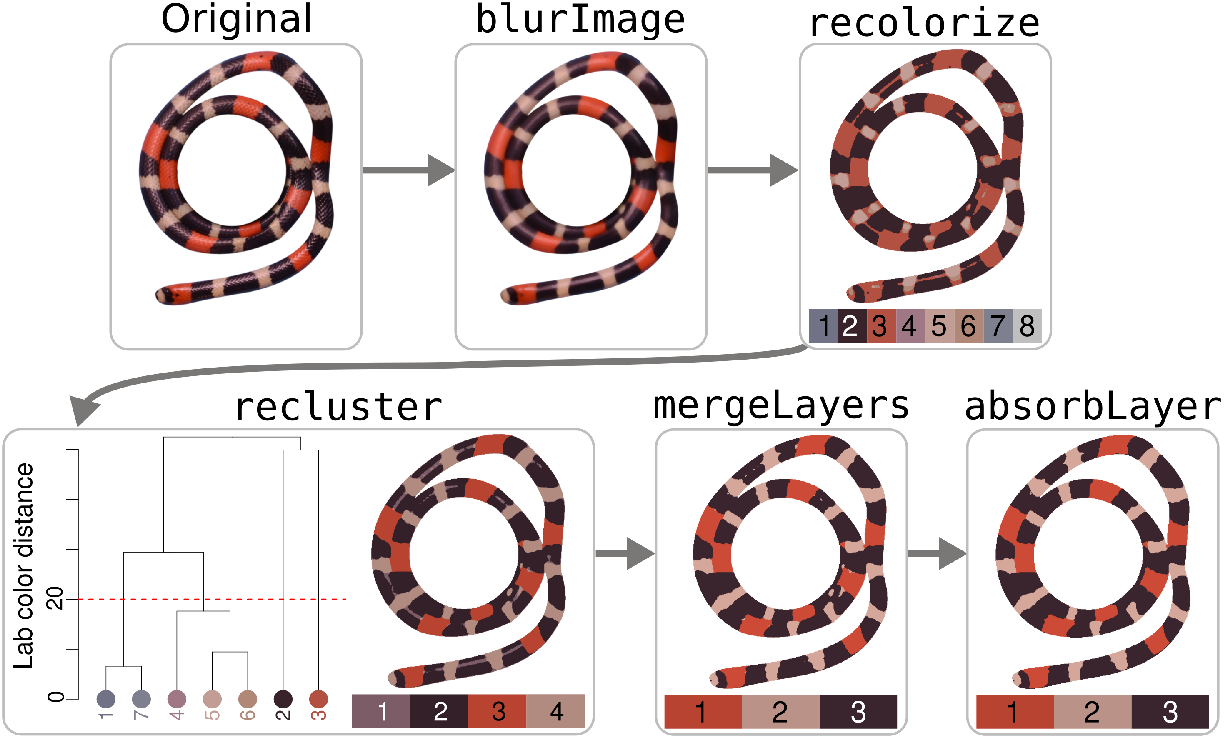
Successful segmentation of the coral snake (*Micrurus* sp.) image from Fig. 2B. This image requires more steps to deal with color variation due to scales and specular highlights: first we blur the image using `blurImage` to mitigate the scale texture, then call `recolorize` (2 bins per channel = 2^3^ = 8 color centers) and `recluster` (cutoff = 20) to perform segmentation. The `mergeLayers` function allows us to specifically combine color centers 1 and 2 (becoming the new color class 3), which eliminates much but not all of the specular highlights. Finally, the `absorbLayer` function eliminates the remaining specular highlights by absorbing isolated speckles from patch 2 into the surrounding color patches.

#### Example C: Lighting variation

Although the ideal image set is acquired using the same camera, lighting conditions, and color standards, users will often have images accumulated from a range of sources where these variables cannot be controlled, e.g. images taken from the iNaturalist database (Nugent, 2018) or by different lab members over several years of a field season. Although the actual color of the image cannot be used for analysis, these images still contain information: we can measure the spatial distribution of colors to quantify color pattern variation, e.g. with `patternize` (Van Belleghem et al., 2018).

Fig. 2C provides a good example of this problem: we know that these images contain identical color pattern information (since this is the same specimen), but the colors themselves vary from a purplish-blue under diffuse light to black with intense specular highlights under direct light. Although k-means clustering produces a dark and a light cluster for each image, the color centers themselves are not only quite different, but the color patches are very different in shape due to the highlights. In this case, we can use `recolorize` (following a similar procedure to that in Fig. 4) to correct for that difference and produce two near-identical color maps from the two images, recovering the same spatial information for the distribution of the light and the dark colors (Fig. 5). Although even in this case the two final maps are not perfectly identical—the borders between the light and dark patches differ slightly (Fig. 5, bottom right)—only 3,270 of 219,107, about 1.5%, of the pixels are assigned differently in each image.

**Figure 5:**
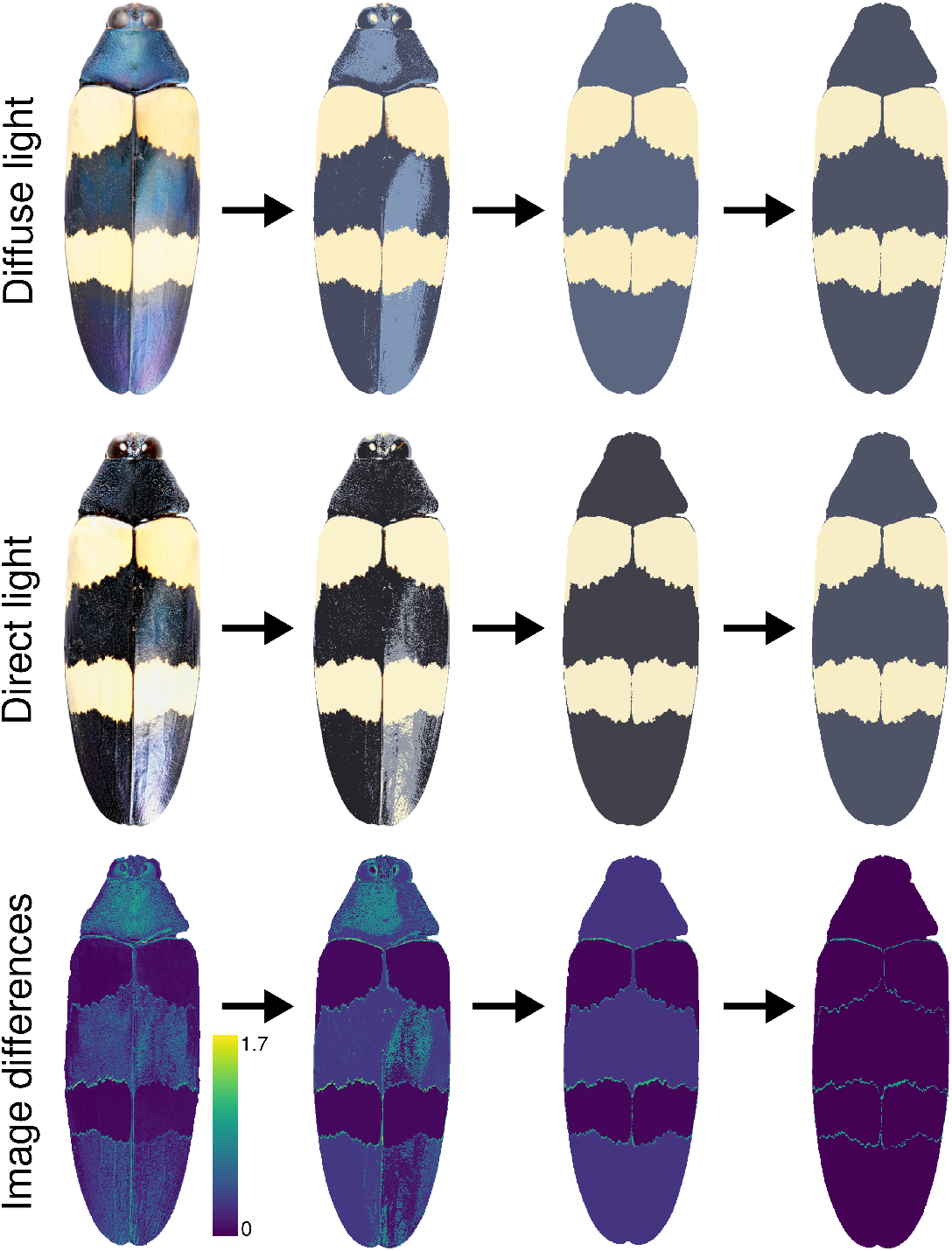
Using `recolorize` to recover the same color pattern information from photos with different lighting conditions. Top row: steps for producing a color map for the diffuse light image. Middle row: steps for the direct light image. Bottom row: differences between the images (calculated using the `imDist` function).

#### Example D: Pattern analysis with patternize

Users will often need to map each image in a dataset to the same color palette in order to compare differences in color pattern distribution. The `patternize` package (Van Belleghem et al., 2018) implements this approach: images are aligned to a common sampling grid (a `RasterStack`) using image registration or landmarks, mapped to a common color palette, and then analyzed with principal components analysis (PCA). This method is probably the closest available equivalent to geometric morphometrics with color pattern analysis, because the alignment step allows us to control for variation in shape, orientation, and size before analyzing variation in color pattern.

In practice, the most difficult step of using `patternize` is often the color segmentation. Like several other color analysis packages and tools, `patternize` mostly relies on k-means clustering to extract color patches from images (especially for batch processing), which can fail for any of the reasons outlined above. Instead, we can combine the strengths of `patternize` and `recolorize` by using `patternize` to perform the image alignment step and `recolorize` to do the color segmentation. Here we show a working example using Polistes fuscatus wasp face images, a subset used with permission from Tumulty et al. (2021). We landmarked the original images (Fig. 6A) in ImageJ, using a simple scheme of only 8 landmarks, along with masking polygons to restrict our analyses to the frons and clypeus of the head (Fig. 6B). These were passed through the `alignLan` function in `patternize` to produce a list of aligned `RasterBrick` objects. We converted these to image arrays using the `brick_to_array` function in `recolorize`, determined a universal color palette using an initial color segmentation of all images (Fig. 5C), then used the `imposeColors` function to map all aligned images to the same set of colors (Fig. 5D). Finally, we converted these color maps back to `patternize` rasters using the `recolorize_to_patternize` function and used the `patPCA` function in `patternize` to perform a whole color pattern PCA (Fig. 5E). A complete version of this example—including step-by-step code to reproduce it and more detailed explanations—is available here: https://hiweller.rbind.io/post/recolorize-patternize-workflow/.

**Figure 6:**
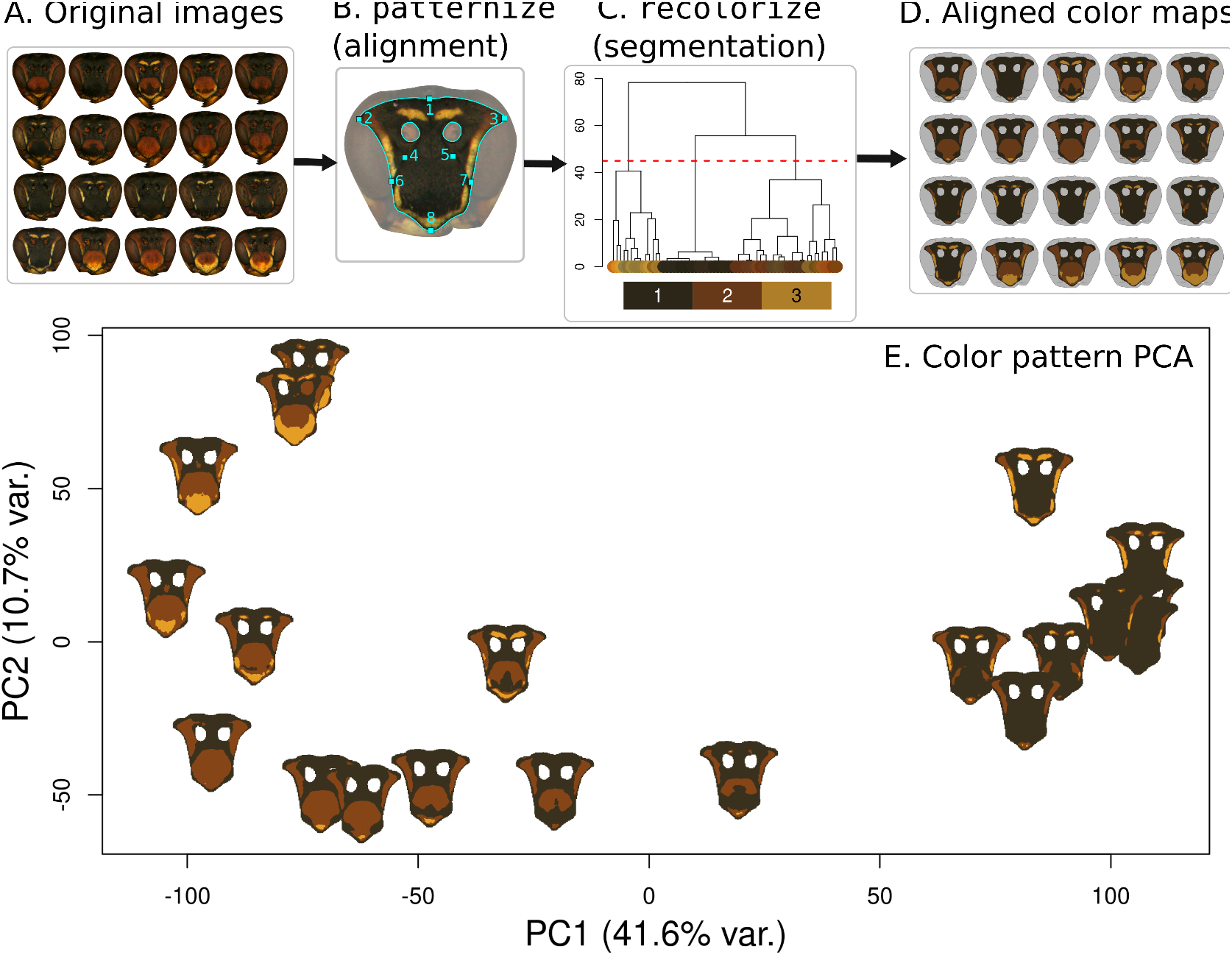
Process for combining `recolorize` and `patternize` to run a whole-colorpattern principal components analysis (PCA). A: Original images of *Polistes fuscatus* wasp faces, taken with the same lighting and camera conditions. B: Landmarking scheme for alignment. C: Generating a universal color palette from color segmentation on individual images. D: Applying the universal color palette using the `imposeColors` function to generate color maps. E: Color pattern PCA as generated by aligned, segmented images, characterizing color pattern diversity among the wasp face images.

#### Example E: Spectral analysis with pavo

Even under ideal circumstances (identical lighting and camera setups, color correction, camera calibration, etc), color images do not provide full-spectrum color information. Instead, they are limited to the visible spectrum and biased toward human visual sensitivities—any user interested in non-human visual perception cannot rely on traditional RGB image data alone to test their questions. One relatively cheap and accessible solution to this problem is to combine color maps, which provide spatial information, with reflectance measurements taken with a point spectrometer.

Reflectance spectra provide a more objective measurement of color and capture wavelengths outside the range of human perception (ultraviolet, 300-400 nanometers) without the expense of obtaining a UV photography set-up. If users have reflectance spectra, the color standardization in their images is less important (so long as spectra can still be accurately assigned to the corresponding color patch). For example, if photos and reflectance spectra were obtained from the same individuals but under variable lighting conditions or camera setups, reflectance spectra could be used to retroactively apply color-standardized data to the images. But because they are point measurements, spectral data cannot provide spatial information. This is part of the workflow of the `pavo` package (Maia et al., 2019), which enables users to combine spatial and spectral information by providing visual models (based on reflectance spectra) in combination with color maps. As with `patternize`, the `pavo` package relies primarily on k-means clustering to do the color segmentation, meaning users encounter many of the pitfalls illustrated in earlier examples.

Here, we illustrate the combination of reflectance spectra with `recolorize` using an example dataset of birds, flowerpiercers in the genus Diglossa (Fig. 7A), which have striking examples of color pattern repetition within and between species (Remsen Jr, 1984; Vuilleumier, 1969) and colors outside of the human visual spectrum. Birds exhibit nearly all of the features which we have shown tend to foil k-mean clustering (Mason et al., 2021). The most common type of bird specimen preparation (aptly termed a ‘round skin’) is convex, with contours that photograph as color variation similar to Fig. 2A; different arrangements of feathers create textures and irregular shadows and shine as in Fig. 2B; and many bird species have finescale color pattern elements like speckles, wingbars, or facial markings which will usually be absorbed by larger color patches. These features make it difficult to combine reflectance spectra with useful color maps.

**Figure 7:**
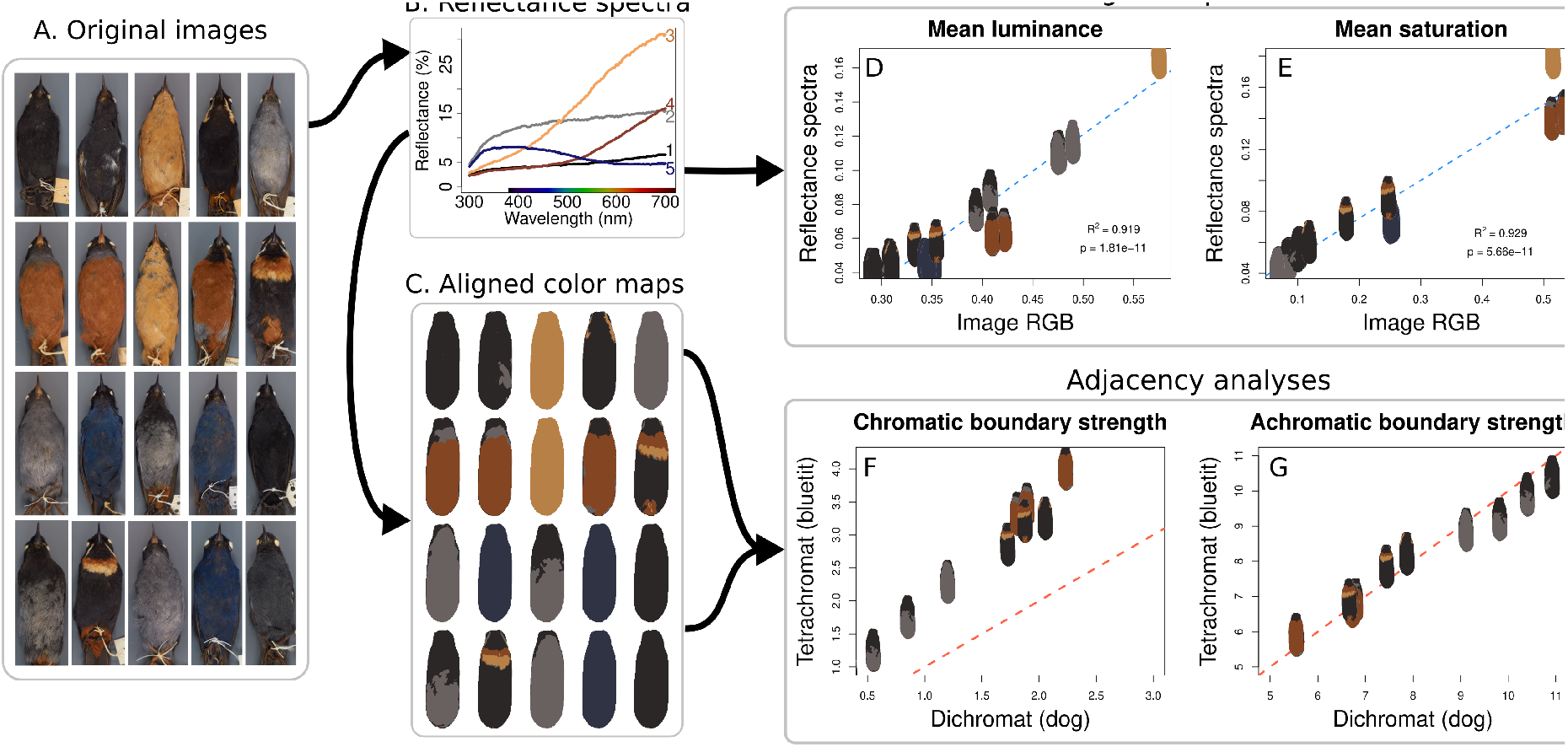
Combining color maps with spectral data in `pavo`. A: Original images of Diglossa spp. birds. B: Reflectance spectra used to determine the five color classes for color segmentation (black, gray, yellow, brown, and blue). C: Color maps (generated in a similar manner to Fig. 6) mapping each bird breast to one of the five color classes. D-E: Comparison of mean luminance and saturation results using image data and spectral data. Blue lines are linear regression fits. F-G: Comparisons of Endler’s adjacency analysis as performed in `pavo` for a dichromat and a tetrachromat (Maia et al., 2019; Endler, 2012). The red dashed line represents a slope of 1 and intercept of 0; points on this line would indicate identical values for the two visual systems. Chromatic boundary strength = color contrast, achromatic boundary strength = luminance (brightness) contrast. Specimens from left to right: row 1 LSUMZ 85389, LSUMZ 90468, LSUMZ 91068, LSUMZ 98863, LSUMZ 98873; row 2 LSUMZ 33373, LSUMZ 33374, LSUMZ 38886, LSUMZ 79148, LSUMZ 80877; row 3 LSUMZ 189657, LSUMZ 196409, LSUMZ 228146, LSUMZ 229100, LSUMZ 229115; row 4 LSUMZ 125407, LSUMZ 129286, LSUMZ 163814, LSUMZ 174295, LSUMZ 179214.

We followed the same procedure as in example D to generate the color maps for the birds. To identify a universal color palette to which bird images should be mapped, we measured reflectance spectra in five locations on the breast of each bird and grouped spectra with similar shapes as calculated using the `peakshape` function in `pavo` (Fig. 7B). In this case, because the reflectance spectra indicated that the navy blue color (color center 5) had higher UV reflectance than the black (color center 1), we kept these as separate color centers although they were very similar in RGB color—an example of using outside information to inform our choice of color palette. We processed bird images through `patternize` and `recolorize` as in Fig. 5 to control for shape variation, then converted the color maps to classify objects using `recolorize_classify` (Fig. 7C).

To combine these color maps with spectral data, we first calculated aggregate reflectance spectra using `procspec` and `aggspecec` in `pavo`, resulting in 5 reflectance curves corresponding to the 5 color centers in the color maps. We used this reflectance data to generate visual models and color distance objects using the `vismodel` and `coldist` functions from the `pavo` package (Maia et al., 2019), here focusing on visual models for a UV-sensitive tetrachromat (bluetit, as provided by `pavo`) and a dichromat (dog).

We then combined spectral and spatial data by running the `adjacent` function in `pavo`, which performs Endler’s adjacency analyses, using these visual models and our `recolorize`-generated color maps. For comparison, we also ran adjacency analysis using the intrinsic RGB colors of the actual color maps, rather than spectral data, which is a simplifying assumption made by the `recolorize_adjacency` function in the absence of spectral data.

We found that the mean luminance and saturation calculated using the intrinsic RGB colors were tightly correlated with these values as calculated using reflectance spectra (R2 = 0.919 and 0.929, respectively), although the scales of these values were quite different for the two methods (Fig. 7D-E). When comparing between dichromatic and tetrachromatic visual systems, we found that the chromatic (color contrast) boundary strength scores differed substantially, with the tetrachromatic visual system having universally higher chromatic boundary strength scores. Interestingly, the achromatic (brightness contrast) boundary strength scores were nearly identical for the two visual systems (Fig. 7F-G).

This example is undoubtedly the most complex of those we present here. We used `patternize`, `recolorize`, and `pavo` to combine spatial and spectral data, analyzing images which pose many of the problems that traditional segmentation methods cannot resolve, and calculating biologically relevant metrics for two different visual models. `Recolorize` worked well for classifying and grouping color patches both within and between species, without the loss of the fine scale pattern information (e.g., the chest bands and mustaches), thus addressing many of the ‘signal-to-noise’ problems that have been a bottleneck for color pattern analysis in birds. Additionally, by integrating `recolorize` with `pavo` we were able to successfully combine spectral data with visual photographs to get a more accurate representation of colors with reflectance in the UV range, a critical component to quantifying color data for taxa like birds which detect colors outside of the human visual range.

#### Example F: Batch processing with different colors

Our final example illustrates an aspect of the `recolorize` workflow which works well for batch processing an image set that does not have a shared color palette. When dealing with a set of images where not each image can be mapped to the same set of colors–for example, a comparative dataset consisting of images of different species–researchers must either fit the same number of color centers to each image, resulting in over- and underclustered images (see Fig. 2D) or choose a different number of colors for each individual image, meaning they have to invent some criterion for determining how many colors to assign each image. In practice, these criteria tend to be fairly subjective. Because the automatic `recolorize` functions operate by grouping colors together by similarity, applying the same series of `recolorize` calls to each image produces a different number of color centers depending on the image (Fig. 8).

**Figure 8:**
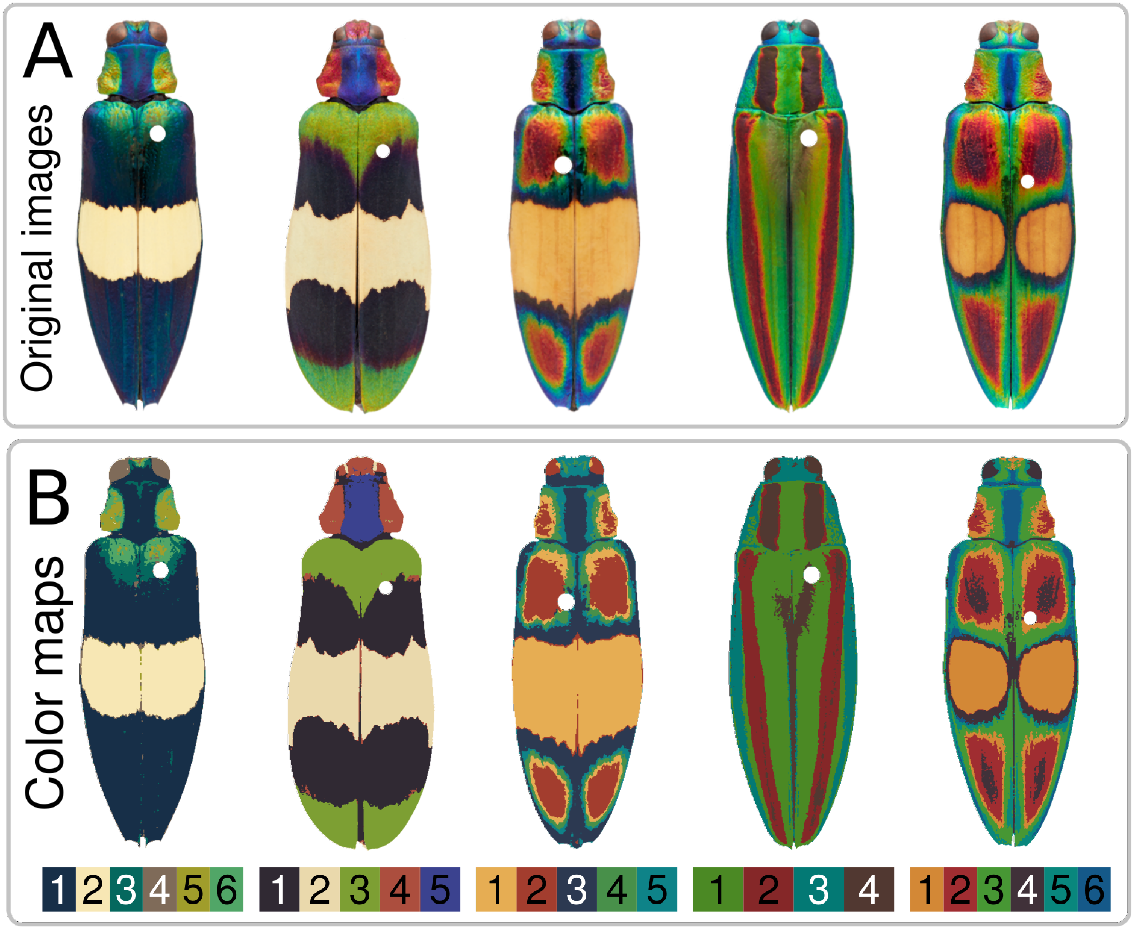
Color maps for *Chrysochroa* spp. beetles, generated by the same series of `recolorize` calls for different images. A: Original images. The white spot on each beetle is where the insect pin was masked with transparency before loading the images into the package. B: Resulting color maps, which range from 4 to 6 colors for each image in this case. Image sources: Nathan P. Lord.

These color maps are imperfect: each beetle in the dataset has a different relationship between texture, shine, and color, which cannot easily be automated in the same call. The initial `recolorize` call could be used to determine the color classes for each image, but users should still go through color maps and make individual modifications as-needed.

### Comparison with existing methods

Although k-means clustering is the most widely used method for color clustering in images, here we compare recolorize to a number of other methods that researchers might encounter when searching for color segmentation solutions (Fig. 9). We summarize the major differences in color clustering methods discussed in this paper in Table 1. In comparison to the other methods, because recolorize includes tools for modifying output that is close to satisfactory, users do not need to find a single solution that will perfectly segment all of their images; they can modify output on a per-image basis, the steps of which are all recorded in the recolorize object for repeatability (see below section on object structure).

**Figure 9:**
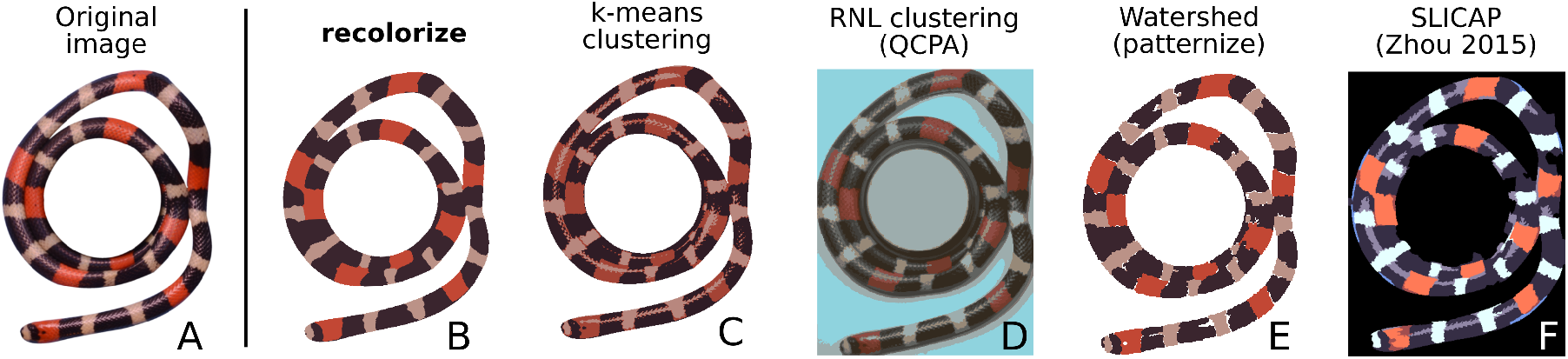
Comparison of recolorize with three other methods for organismal color segmentation. A: Original image. B: Recolorize output as achieved in Fig. 4. C: One run of k-means clustering output. D: Output from receptor noise-limited (RNL) clustering as implemented by the QCPA framework in micaToolbox (van den Berg et al., 2020; Troscianko and Stevens, 2015). E: Watershed segmentation of three colors in patternize (separate layers are superimposed). F: Simple linear iterative clustering with affinity propagation (SLICAP) segmentation as described by Zhou (2015) and implemented by Lampros (2021)

#### Receptor noise-limited clustering

The ImageJ plugin micaToolbox (Troscianko and Stevens, 2015) and accompanying Quantitative Color Pattern Analysis toolkit (van den Berg et al., 2020) are among the most comprehensive tools widely available to biologists for modeling non-human visual systems. One option in the toolkit is receptor noise-limited (RNL) color clustering, which uses perceptual thresholds of a specified visual system to cluster an image based on whether a given viewer could distinguish colors at a specified viewing distance. To run RNL clustering, we used a camera RAW image of the snake that included a 40% reflectance standard, as well as calibrating the camera using a separate image of an Xrite Colorchecker. We then generated a multispectral image, used region-of-interest (ROI) masking to analyze only the snake, performed acuity correction for a viewing distance of one meter, converted to a cone-catch image (we chose a human model for comparison with other methods), ran the RNL ranked filter, and finally RNL clustering (van den Berg et al., 2020; Caves and Johnsen, 2018). The resulting image includes some background (despite the ROI implementation) and segments the snake itself into 19 color clusters. With ROI masking, processing this image took three minutes on a personal laptop with 16Gb RAM (not accounting for user error); without ROI masking, the image took over 20 minutes to process.

#### Watershedding in patternize

The watershedding algorithm as implemented in patternize is intended to solve problems of shine and texture on an image. Although the watershed output (as implemented by the patLanW function in patternize) is usable, it requires repeated user input (clicking on the original image to set seeds for each discrete color patch, e.g. every black segment of the snake) for each color cluster, and in this case the results still do not completely solve the specular reflectance problem (Fig. 9D), meaning in this case the method is both less effective and more subjective.

#### General color segmentation algorithm

We also attempted to use an algorithm for general color segmentation of images as described by Zhou (2015), termed simple linear iterative clustering with affinity propagation (SLICAP). Given that this method also does not require any a priori specification of the expected number of colors, it performs remarkably well (producing 6 color clusters not counting the background), but still results in many color clusters with no easy way for users to modify the output.

### Package installation, structure, and input

#### Installation

The most recent stable release version of the package can be installed from the Comprehensive R Archive Network (CRAN) from R using the `install_packages()` function:

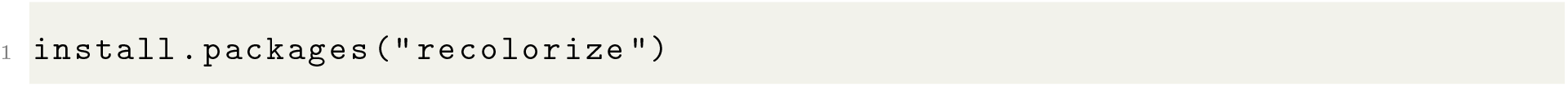

The development version of the package can be installed from GitHub (https://github.com/hiweller/recolorize) using the devtools package (Wickham et al., 2021):

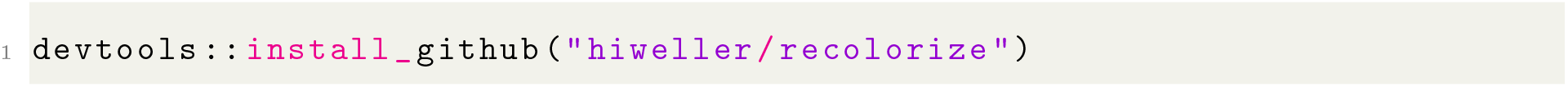

### The recolorize class

The `recolorize` package mostly works with R objects of S3 class `recolorize`, which are output by the base functions and which most functions in later steps of the workflow will take in as an argument. Objects of this class are lists with the following elements:

1. `original_img:` The original image, stored as a raster array (essentially a matrix of hexadecimal codes).
2. `centers:` A matrix of color centers, listed as one RGB triplet per row in a 0-1 range. These are usually the average color of all pixels assigned to that color class unless otherwise specified by the user.
3. `sizes:` The number of pixels assigned to each color class.
4. `pixel_assignments:` A matrix of color class assignments for each pixel. For example, all pixels coded as 1 in the `pixel_assignments` matrix are assigned to color class 1 (which will be row 1 of `centers`).
5. `call:` The set of commands that were called to generate the `recolorize` object.

The call is especially helpful for reproducibility, because it stores every step used to generate the current segmentation (any function that returns a `recolorize` class object will modify the call element accordingly).

## Discussion

Fully automated methods rarely work all of the time, and are difficult to modify, while fully manual methods are subjective and time-consuming. `Recolorize` strikes a balance between these two extremes by providing an effective color segmentation algorithm along with tools for modifying and exporting the resulting color maps. In the simplest case, users only have to tinker with the number of initial color centers in the first step and the similarity cutoff in the second step. Even in more complicated cases, where color maps are modified individually, these steps are recorded in the call element of the `recolorize` objects. This design allows `recolorize` to handle a much wider range of color segmentation problems than it could otherwise.

|  | recolorize | k-means<br>clustering | RNL<br>clustering | Watershed | SLICAP |
| --- | --- | --- | --- | --- | --- |
| Images from multiple sources | Y | Y |  | Y | Y |
| No calibration required | Y |  |  | Y | Y |
| Automated clustering options | Y |  | Y |  | Y |
| Tools for modifying output | Y |  |  |  |  |
| Deterministic | Y |  | Y |  | Y |
| Batch processing (diff. colors) | Y | Y | Y |  | Y |
| Batch processing (same colors) | Y | + |  | Y |  |
| Export directly to other methods | Y |  |  |  |  |
| Supports transparencies | Y |  |  |  | + |
| Multispectral images |  |  | Y |  |  |
| Non-human visual systems |  |  | Y |  |  |
| Graphical user interface |  |  | Y | Y |  |

### Comparison and complementarity with existing methods

Most of the methods to which we compared recolorize are implemented in existing pipelines, and are therefore not the sole focus of the software or method in question. For example, the RNL clustering output in Fig. 9C requires specific calibration equipment, knowledge of visual system parameters, and ten processing steps to achieve a 19-cluster color map. The results are accurate to human perception: portions of the snake are different colors due to shine and 3D contour, which would be detectable to a human viewer. However, the extra equipment and number of steps required for processing are prohibitive given the research context. This would restrict the analysis to images taken using the same camera and which include a relatively expensive calibration standard, and while the results are highly informative for visual perception, users would have to do substantial modification to measure, for example, the proportion of black on the snake’s body, or the length of the border shared between the red and white patches, since each of these is broken up into multiple clusters which sometimes span more than one color. This is because the QCPA workflow in general is concerned with simulating non-human visual systems, which requires a higher standard of calibration and more carefully controlled data collection.

Because recolorize is a dedicated toolbox for organismal color segmentation, it is designed not as a replacement for existing pipelines but as a complement to them. By making color segmentation more feasible and providing export options to a variety of formats for multiple user cases, recolorize makes other color analysis tools easier to use for a wider variety of projects and images.

### Current and potential applications for recolorize

Currently, `recolorize` works with PNG and JPEG images, and does not support less common (but more information-rich) formats, such as the multispectral images generated with micaToolbox (van den Berg et al., 2020) or Image Calibration Analysis Toolbox (Troscianko and Stevens, 2015). However, the underlying package structure can be extended to other formats as the central algorithms of the package can be modified to images with more than 3 channels, and intermediate steps are exported as their own functions (in addition to being called on by `recolorize()`). For example, we recently used `recolorize` functions for color segmentation of 3D objects (STL files output from photogrammetry; Christopher Taylor, pers. comm.). Such future developments, often driven by specific user cases, will be made available on GitHub.

The `recolorize` toolbox can be used to process a high number of images more consistently than existing manual or simple automated methods, but its output is imperfect. Users are invariably going to have to tweak problem images or do some things manually if they want 100% efficacy, and will otherwise have to accept some amount of error. In some ways this is about choosing your source of error: computer or user?

The relative ease with which we can combine color maps with spectral data (per example E) also suggests interesting possibilities. Even in this reduced example, when we compare chromatic and achromatic boundary strength for the tetrachromat and dichromat, we see that chromatic boundary strength (color contrast) is measurably different between the two visual systems (generally higher for the tetrachromat than the dichromat), which we would expect. However, we also see that the two visual systems are very closely matched in achromatic boundary strength (brightness contrast), which suggests that achromatic boundary strength depends less on particular properties of a given visual system than chromatic boundary strength. When we measured mean luminance and saturation from reflectance spectra versus intrinsic RGB colors, we found tight correlations (but different scales) for the two sets of measurements. In this case, because the Diglossa dataset contains high-quality images acquired under consistent settings, it would not be wildly unreasonable to use the intrinsic RGB colors if spectral data were unavailable. This approach must be used with caution, especially if researchers know of (or are uncertain about) a substantial UV-reflective component of the color patterns in question, or if the images are from different sources.

A last possibility would be to use `recolorize` to generate a training set for a machine learning approach. These generally have the problem, especially in fields like organismal biology, that the amount of training data and expertise required to get a sufficiently trained algorithm is actually more effort than just doing everything manually (given that it usually has limited applicability). Performing the segmentation in `recolorize` might make it easier to generate that training data so this solution could be used for more specific problems.

### The ‘correct’ color map depends on the question

Image segmentation is a classically difficult problem in computer vision, especially because there is no single ‘correct’ answer for appropriate color pattern segmentation. A color map is by definition a simplified representation of an actual color pattern, so the correct solution is not intrinsic to the image, but depends on the user’s question. For example, in Fig. 10, we illustrate two possible color segmentations for our original image of an iridescent jewel beetle (*Chrysochroa fulgidissima*). In Fig. 10B, we show a more complex, 8-color segmentation, which retains the brighter orange on the borders of the red stripes, and fits different shades of red and blue to reflect differences in viewing angle of the iridescent elytra. This color map would be appropriate for answering questions of visual contrast and perception, since it retains more properties relevant for visual stimuli. In Fig. 10C, we show a much simpler 2-color segmentation, consisting only of red and green. This is not a very visually faithful representation of the original image, but if we wanted to measure the location and distribution of green iridescence across beetle taxa, this map would be much more helpful to us than that in Fig. 10B. We end on this example to emphasize that there is no universal solution for the problem of biological color segmentation: there is no method so comprehensive that it absolves researchers of posing specific questions.

**Figure 10:**
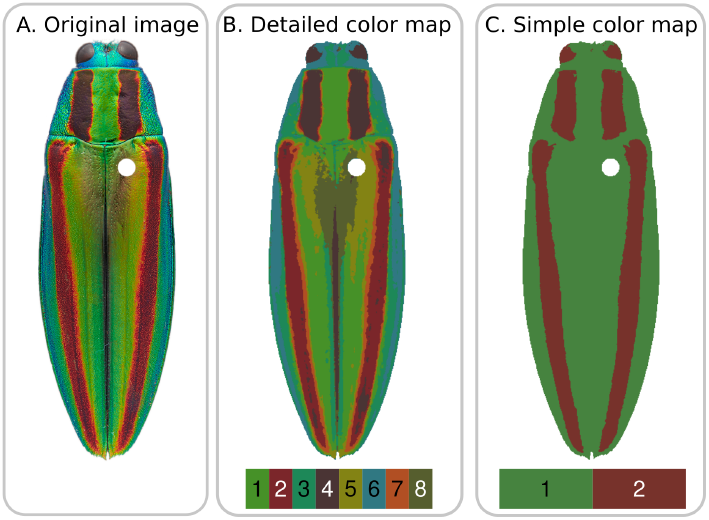
Different color maps generated for the same image. Solution depends on the use case. Image: Nathan P. Lord.

## Acknowledgements

We are grateful to James Tumulty, Alison Davis-Rabosky, and Able Chow for providing images used as examples. Christopher Taylor, Ghislaine Cardenas-Posada, Matthew Fuxjager, John Capano, and Nicole Moody provided helpful feedback on the first draft of the paper. Thank you to the Louisiana State University (LSU) color seminar group and Brant Faircloth for sparking the discussions that led to this paper. Thanks to Robb Brumfield, Nick Mason, and the LSU Museum of Natural Science for access to and resources for photographing bird specimens. Lastly, we would like to thank HIW’s PhD advisor, Elizabeth Brainerd, for providing HIW with the academic freedom and gentle but needed encouragement to pursue this project.

## Attributions

HIW wrote the package, example code, and manuscript. NPL provided images, feedback, funding, and helped write MS. AEH provided bird data (images and spectra) and helped write the manuscript. SMVB helped extend the package to be compatible with `patternize` and helped with coding, especially in the wasp example, and edited the manuscript.

## Data availability

The development version of the package is available on GitHub: https://github.com/hiweller/recolorize. The stable release version is available on the Central R Archive Network: https://cran.r-project.org/package=recolorize. All images and code used to generate the examples are available in a separate Github repository: https://github.com/hiweller/recolorize_examples.

## References

Dean C. Adams and Erik Otárola-Castillo. geomorph: an r package for the collection and analysis of geometric morphometric shape data. Methods in Ecology and Evolution, 4(4):393–399, 2013. doi: https://doi.org/10.1111/2041-210X.12035. URL https://besjournals.onlinelibrary.wiley.com/doi/abs/10.1111/2041-210X.12035.

Dean C Adams, F James Rohlf, and Dennis E Slice. Geometric morphometrics: ten years of progress following the ‘revolution’. Italian Journal of Zoology, 71(1):5–16, 2004.

H. W. Bates. A Naturalist on the River Amazons. John Murray, 1863.

I. Bekker, F. Sylburg, and K.F. Neumann. *Historia animalium*. Aristotelis Opera. e Typographeo academico, 1837. URL https://books.google.com/books?id=2Z8NAAAAYAAJ.

Fred L Bookstein. Combining the tools of geometric morphometrics. In Advances in morphometrics, pages 131–151. Springer, 1996.

Eleanor M Caves and Sönke Johnsen. Acuityview: An r package for portraying the effects of visual acuity on scenes observed by an animal. Methods in Ecology and Evolution, 9(3):793–797, 2018.

Ian ZW Chan, Martin Stevens, and Peter A Todd. Pat-geom: a software package for the analysis of animal patterns. Methods in Ecology and Evolution, 10(4):591–600, 2019.

John David Curlis, Timothy Renney, Alison R Davis Rabosky, and Talia Y Moore. Batch-mask: An automated mask r-cnn workflow to isolate non-standard biological specimens for color pattern analysis. bioRxiv, 2021.

John A. Endler. A framework for analysing colour pattern geometry: Adjacent colours. Biological Journal of the Linnean Society, 107(2):233–253, 2012. ISSN 00244066. doi: 10.1111/j.1095-8312.2012.01937.x.

John A. Endler, Gemma L. Cole, and Alexandrea M. Kranz. Boundary strength analysis: Combining colour pattern geometry and coloured patch visual properties for use in predicting behaviour and fitness. Methods in Ecology and Evolution, 9(12):2334–2348, 2018. ISSN 2041210X. doi: 10.1111/2041-210X.13073.

Briana D Ezray, Drew C Wham, Carrie E Hill, and Heather M Hines. Unsupervised machine learning reveals mimicry complexes in bumblebees occur along a perceptual continuum. Proceedings of the Royal Society B, 286(1910):20191501, 2019.

Felipe M. Gawryszewski. Color vision models: Some simulations, a general n-dimensional model, and the colourvision r package. Ecology and Evolution, 8(16):8159–8170, 2018. doi: https://doi.org/10.1002/ece3.4288. URL https://onlinelibrary.wiley.com/doi/abs/10.1002/ece3.4288.

John A Hartigan and Manchek A Wong. Algorithm as 136: A k-means clustering algorithm. Journal of the royal statistical society. series c (applied statistics), 28(1):100–108, 1979.

Sarah E Hooper, Hannah Weller, and Sybill K Amelon. Countcolors, an r package for quantification of the fluorescence emitted by pseudogymnoascus destructans lesions on the wing membranes of hibernating bats. Journal of Wildlife Diseases, 56(4):759–767, 2020.

Sönke Johnsen. The optics of life. Princeton University Press, 2012.

Christian Peter Klingenberg. Morphoj: an integrated software package for geometric morphometrics. Molecular ecology resources, 11(2):353–357, 2011.

Mouselimis Lampros. SuperpixelImageSegmentation: Image Segmentation using Superpixels, Affinity Propagation and Kmeans Clustering, 2021. URL https://CRAN.R-project.org/package=SuperpixelImageSegmentation.

A Michelle Lawing and P David Polly. Geometric morphometrics: recent applications to the study of evolution and development. Journal of Zoology, 280(1):1–7, 2010.

Rafael Maia, Hugo Gruson, John A Endler, and Thomas E White. pavo 2: new tools for the spectral and spatial analysis of colour in r. Methods in Ecology and Evolution, 10(7):1097–1107, 2019.

Nicholas A Mason, Eric A Riddell, Felisha Romero, Carla Cicero, and Rauri CK Bowie. Plumage balances camouflage and thermoregulation in horned larks (eremophila alpestris). BioRxiv, 2021.

Jill Nugent. inaturalist. Science Scope, 41(7):12–13, 2018.

Aaron M Olsen and Mark W Westneat. Stereomorph: An r package for the collection of 3d landmarks and curves using a stereo camera set-up. Methods in Ecology and Evolution, 6(3):351–356, 2015.

Anna Orteu and Chris D. Jiggins. The genomics of coloration provides insights into adaptive evolution. Nature Reviews Genetics, 21(8):461–475, 2020. ISSN 14710064. doi: 10.1038/s41576-020-0234-z. URL http://dx.doi.org/10.1038/s41576-020-0234-z.

P David Polly, A Michelle Lawing, Anne-Claire Fabre, and Anjali Goswami. Phylogenetic principal components analysis and geometric morphometrics. Hystrix, 24(1):33, 2013.

E.B. Poulton. The Colours of Animals: Their Meaning and Use, Especially Considered in the Case of Insects. International scientific series. D. Appleton, 1890. URL https://books.google.com/books?id=Uf4vAAAAYAAJ.

R Core Team. R: A Language and Environment for Statistical Computing. R Foundation for Statistical Computing, Vienna, Austria, 2022. URL https://www.R-project.org/.

JV Remsen Jr. High incidence of ”leapfrog” pattern of geographic variation in andean birds: implications for the speciation process. Science, 224(4645):171–173, 1984.

Shawn T Schwartz and Michael E Alfaro. Sashimi: A toolkit for facilitating high-throughput organismal image segmentation using deep learning. Methods in Ecology and Evolution, 12(12):2341–2354, 2021.

Jolyon Troscianko and Martin Stevens. Image calibration and analysis toolbox–a free software suite for objectively measuring reflectance, colour and pattern. Methods in Ecology and Evolution, 6(11):1320–1331, 2015.

Jolyon Troscianko, John Skelhorn, and Martin Stevens. Quantifying camouflage: how to predict detectability from appearance. BMC evolutionary biology, 17(1):1–13, 2017.

James P. Tumulty, Sara E. Miller, Steven M. Van Belleghem, Hannah I. Weller, Christopher M. Jernigan, Sierra Vincent, Regan J. Staudenraus, Andrew W. Legan, Timothy J. Polnaszek, Floria M. K. Uy, Alexander Walton, and Michael J. Sheehan. Evidence for a selective link between cooperation and individual recognition. bioRxiv, 2021. doi: 10.1101/2021.09.07.459327. URL https://www.biorxiv.org/content/early/2021/09/08/2021.09.07.459327.

M Valcu and J Dale. colorzapper: Color extraction utilities. r package version 1.0, 2014.

Jennifer J Valvo, Jose David Aponte, Mitch J Daniel, Kenna Dwinell, Helen Rodd, David Houle, and Kimberly A Hughes. Using delaunay triangulation to sample whole-specimen color from digital images. Ecology and evolution, 11(18):12468–12484, 2021.

Steven M Van Belleghem, Riccardo Papa, Humberto Ortiz-Zuazaga, Frederik Hendrickx, Chris D Jiggins, W Owen McMillan, and Brian A Counterman. patternize: An r package for quantifying colour pattern variation. Methods in Ecology and Evolution, 9(2):390–398, 2018.

Steven M Van Belleghem, Paola A Alicea Roman, Heriberto Carbia Gutierrez, Brian A Counterman, and Riccardo Papa. Perfect mimicry between heliconius butterflies is constrained by genetics and development. Proceedings of the Royal Society B, 287(1931):20201267, 2020.

Cedric P. van den Berg, Jolyon Troscianko, John A. Endler, N. Justin Marshall, and Karen L. Cheney. Quantitative Colour Pattern Analysis (QCPA): A comprehensive framework for the analysis of colour patterns in nature. Methods in Ecology and Evolution, 11(2):316–332, 2020. ISSN 2041210X. doi: 10.1111/2041-210X.13328.

François Vuilleumier. Pleistocene speciation in birds living in the high andes. Nature, 223(5211):1179–1180, 1969.

Hannah I Weller and Mark W Westneat. Quantitative color profiling of digital images with earth mover’s distance using the r package colordistance. PeerJ, 7:e6398, 2019.

Hadley Wickham, Jim Hester, Winston Chang, and Maintainer Jim Hester. Package ‘devtools’. 2021.

Canchao Yang, Longwu Wang, Wei Liang, and Anders P Møller. Do common cuckoos (cuculus canorus) possess an optimal laying behaviour to match their own egg phenotype to that of their oriental reed warbler (acrocephalus orientalis) hosts? Biological Journal of the Linnean Society, 117(3):422–427, 2016.

Bao Zhou. Image segmentation using slic superpixels and affinity propagation clustering. Int. J. Sci. Res, 4(4):1525–1529, 2015.

